# The making of cauliflowers: the story of unsuccessful flowers

**DOI:** 10.1101/2021.02.12.427428

**Authors:** Eugenio Azpeitia, Gabrielle Tichtinsky, Marie Le Masson, Antonio Serrano-Mislata, Veronica Gregis, Carlos Gimenez, Nathanaёl Prunet, Jérémy Lucas, Etienne Farcot, Martin M. Kater, Desmond Bradley, Francisco Madueño, Christophe Godin, Francois Parcy

**Affiliations:** Laboratoire de Reproduction et Développement des Plantes, Univ. Lyon, ENS de Lyon, UCB Lyon 1, CNRS, INRAE, Inria, 46 allée d’Italie, F-69364, Lyon, France; Laboratoire Physiologie Cellulaire et Végétale, Univ. Grenoble Alpes, CNRS, CEA, INRAE, IRIG-DBSCI-LPCV, 17 avenue des martyrs, F-38054, Grenoble, France; Instituto de Biología Molecular y Celular de Plantas (IBMCP), Consejo Superior de Investigaciones Científicas (CSIC) - Universidad Politécnica de Valencia (UPV), 46022 Valencia, Spain; Dipartimento di Bioscienze, Università degli Studi di Milano, Via Celoria 26, 20133 Milan, Italy; Division of Biology and Biological Engineering, California Institute of Technology, 1200 E. California Blvd., Pasadena, CA 91125, USA and Department of Molecular, Cell and Developmental Biology, University of California, Los Angeles, 610 Charles E. Young dr. S., Los Angeles, CA 90095, USA; School of Mathematical Sciences, University of Nottingham, NG7 2RD, United Kingdom; Department of Cell and Developmental Biology, John Innes Centre, NR4 7UH Norwich NR4 7UH, United Kingdom

**Author notes:** Centro de Ciencias Matemáticas, Universidad Nacional Autónoma de México, Morelia, México (Eugenio Azpeitia).

## Abstract

The arrangement of plant organs, called phyllotaxis, produce remarkable spiral or whorled patterns. Cauliflowers present a unique phyllotaxis with a multitude of spirals over a wide range of scales. How such a self-similar fractal organization emerges from developmental mechanisms has remained elusive. Combining experimental assays with modeling, we found that cauliflowers arise due to the hysteresis of the bistable floral network that generates inflorescences imprinted by a transient floral state. We further show how additional mutations affecting meristem growth dynamics can induce the production of conical phyllotactic structures reminiscent of the conspicuous fractal Romanesco shape. This study reveals how the spectacular morphological modification of the inflorescences in cauliflower and Romanesco shape arises from the hysteresis of the genetic programs controlling inflorescence development.

**One Sentence Summary:** The molecular making of cauliflowers

## Main Text

Plant architectures arise from the activity of shoot apical meristems (SAM), which are pools of stem cells fueling the initiation of organs such as leaves, shoots or flowers. The arrangement of organs produced by meristems is named phyllotaxis. In plants with a so-called spiral phyllotaxis, organs usually form two families of spirals that are easily visible when the structure remains compact, such as in flower heads (Fig. 1a,b), pine cones or cactii (Fig. 1c). These two visual families of spirals, called parastichies, turn in opposite directions, and the number of spirals in each direction are usually two consecutive numbers of the Fibonacci series (e.g. 13 blue and 21 red spirals in Fig. 1a) *(1)*. Cauliflowers represent a unique example of spiral phyllotaxis where parastichy family pairs are visible not only at one but at several scales in a massive manner (Fig. 1d-f). This striking self-similar organization culminates in the Romanesco cultivar where the spirals are even more visible due to their conical shape, a geometrical feature conferring the whole curd a conspicuous fractal-like aspect (Fig. 1g).

**Figure 1:**
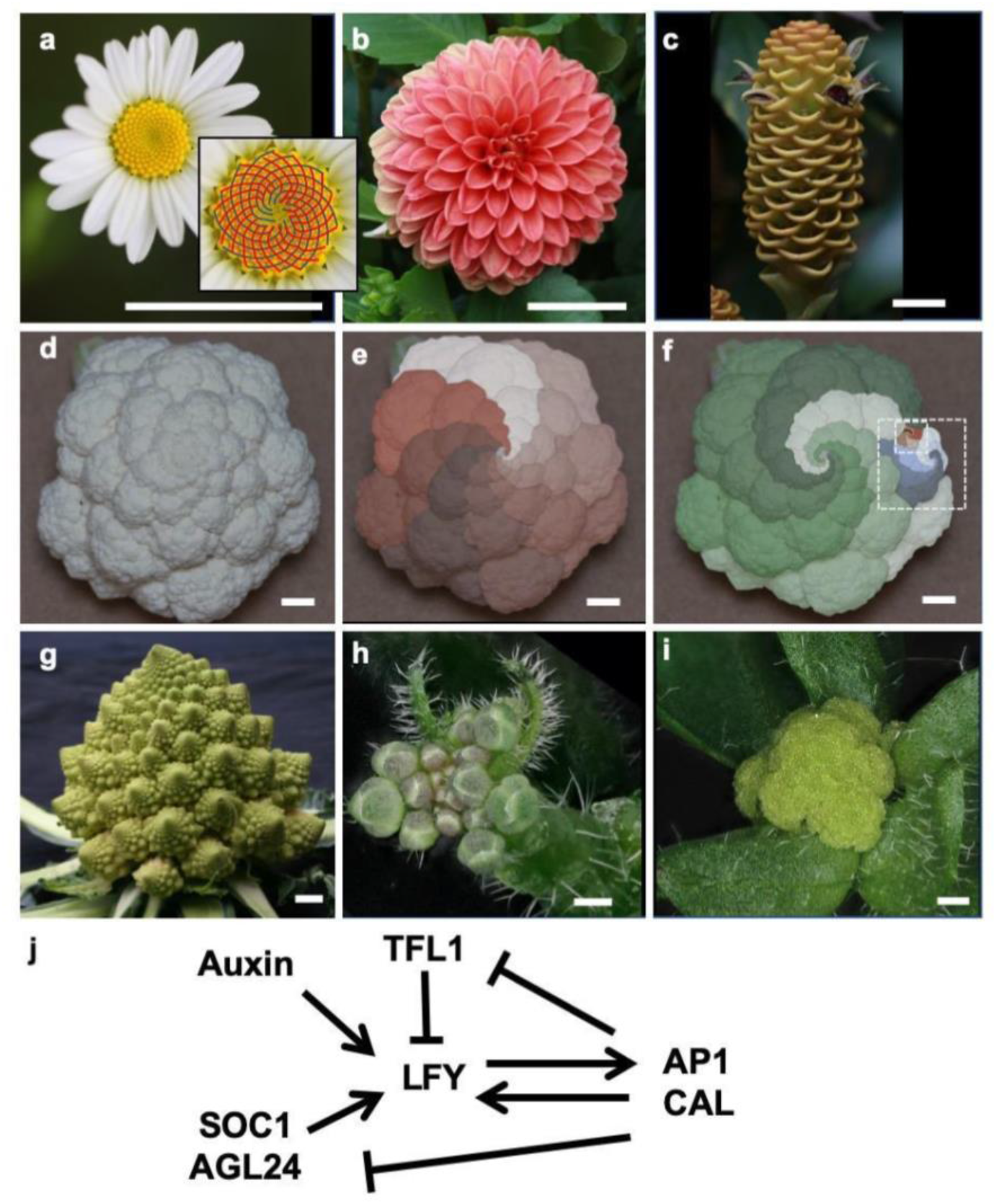
Illustrations of phyllotactic spirals on plant inflorescences. Inflorescences frequently show organs organized in two families of visual spirals turning clockwise and anti-clockwise (parastichies): (a) Daisy capitulum (the two families of spirals are indicated in the close-up by red and blue curves). (b) Dahlia composite flower (c) Zingiber inflorescence. (d-f) *Brassica oleracea* var. *botrytis* cauliflower curd, and (g) *Brassica oleracea* var. *botrytis* ‘Romanesco’ curd. (h) *Arabidopsis thaliana* wild-type flowering shoot, (i) *ap1 cal A. thaliana* curd replacing flowers, Bars = 2 cm for a-g, 500 μm for h-i. (j) Interactions between major floral regulators, arrows depict activation whereas barred lines indicate repression.

Cauliflowers (*Brassica oleracea* var. *botrytis*) were domesticated from the wild *Brassica oleracea (2)*. The cauliflower inflorescence (the flower bearing shoot) takes a curd shape because each emerging flower primordia never fully reaches the floral stage, and repeatedly generates a novel curd-shaped inflorescence instead *(3)*. In *B. oleracea*, the genetic modifications causing curd development are still debated *(4,5)*. However, cauliflowers also exist in the model brassicaceae *Arabidopsis thaliana* and are known to be caused by a double mutation in *APETALA1* (*AP1*) and *CAULIFLOWER* (*CAL*) (Fig. 1h-i), two paralogous genes encoding MADS-box transcription factors (TF) promoting floral development *(3,6)*. The Arabidopsis networks of molecular regulators governing the development of shoots and flowers have been largely identified. However, how variants of those networks *(7–11)* affect growth dynamics so that it would generate the self-nested and compact arrangements of spirals found in cauliflower curds has remained elusive.

To address this question, we first focused on the main molecular actors involved in the floral transition to understand how their network of interactions drives meristem fate. Then, we embedded this molecular network within a 3D computational model of plant development to understand how the molecular changes resulting from the cauliflower mutation may alter organ growth and transform wild-type (WT) inflorescences into spectacular curd phenotypes.

## The genetic basis of cauliflower curds

We first chose to build a minimal regulatory network based on the interactions between the key regulators controlling early floral meristem development *(6,12)*. In Arabidopsis, flowers are initiated by the LEAFY (LFY) transcription factor (Fig. 1j). *LFY* is upregulated by the SUPPRESSOR-OF-OVEREXPRESSION-OF-CO 1 (SOC1) and AGAMOUS-LIKE 24 (AGL24) MADS-box proteins (induced throughout the inflorescence meristem by environmental and endogenous cues) and by the auxin phytohormone maxima that mark floral meristems initiation sites. *LFY* is expressed specifically in floral primordia since its induction in the center of the SAM is repressed by the TFL1 inflorescence identity protein. In the floral primordium, LFY induces *AP1/CAL* that positively feed back on *LFY* and repress *SOC1/AGL24* and *TFL1*, thereby stabilizing the floral fate of the new meristem. In the *ap1 cal* cauliflower mutant, the AP1/LFY positive feedback is absent and *TFL1* is not repressed by AP1/CAL in the nascent floral meristem. Consequently, young flower primordia cannot maintain *LFY* expression and lose their floral identity to become new inflorescence meristems with high *TFL1* expression. Whereas *TFL1* repression in nascent flower primordia is well understood, the factors directly responsible for its upregulation in *ap1/cal* are not known. Their identification is a prerequisite to build a realistic gene regulatory network (GRN) able to drive WT and cauliflower curd development.

To solve this issue, we searched for novel direct positive regulators of *TFL1*. This gene is known to be indirectly regulated by day length *(13)*: in long days (LD) *TFL1* is up-regulated by CONSTANS (CO) and FT, two key upstream effectors of the LD pathway *(7,14–16)* (Fig. S1). To search for direct regulators, we examined SOC1 and AGL24 that act downstream of CO and FT in the LD pathway *(11)*. Loss- and gain-of-function experiments demonstrated that both SOC1 and AGL24 induce *TFL1* (Fig. 2a-i) and Chromatin Immuno-Precipitation experiments showed that these two TFs bind to the *TFL1* regulatory regions previously identified as important for expression in the SAM *(17)* (Fig. 2j-l). These regions were even sufficient to mediate the activation of a *TFL1* reporter construct by SOC1 and AGL24 in a transient assay (Fig. 2m-n) confirming that both MADS-box TFs are direct regulators of *TFL1*. Since XAL2, a close homolog of SOC1 and AGL24 was also shown to bind to and induce *TFL1 (19)*, we aggregated the activities of SOC1, AGL24 and XAL2 into a ‘SAX’ proxy acting as novel *TFL1* positive regulator in our model (Fig. 3a).

**Fig. 2:**
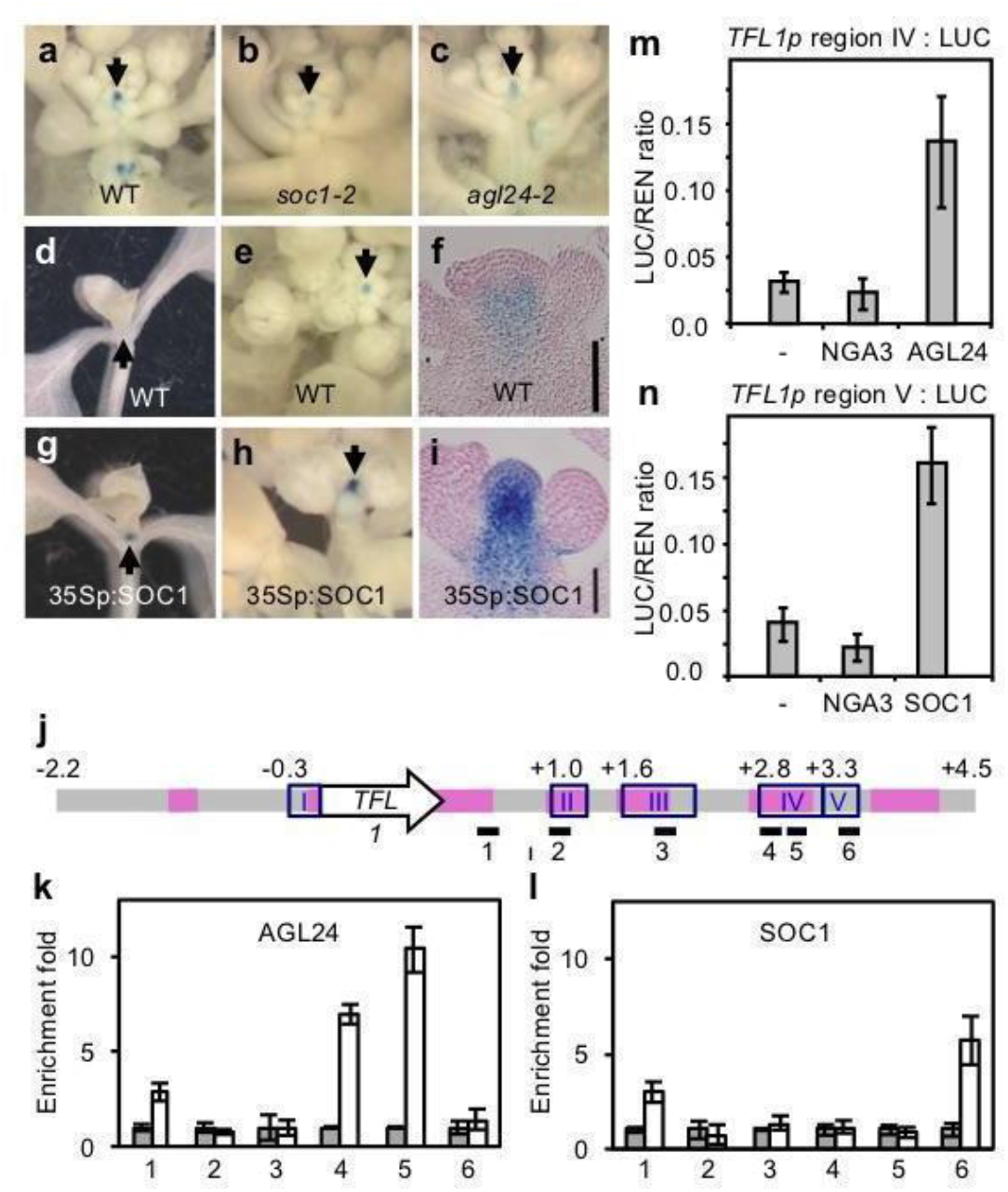
AGL24 and SOC1 are direct positive regulators of *TFL1*. (a-c), TFL1p:GUS activity in WT (a), *soc1-2* (b) and *agl24-2* (c) flowering shoot apices. (d-i), TFL1p:GUS activity in WT (d-f) and *35Sp:SOC1* (g-i) apices at vegetative (d,g) and flowering (e,f,h,i) stages. (f-i), longitudinal sections through flowering shoots. Arrows mark the SAM region. Scale bars in (f) and (i), 40 μm. (j-l) Structure of the *TFL1* genomic region, with regions conserved in Brassicaceae (pink lines), regulatory regions defined in *(17)* (blue boxes I-V), and fragments used for the ChIP with AGL24 and SOC1 (black lines 1-6). ChIP experiments performed on plants expressing a tagged version of AGL24 (k, white bars) or the wild type SOC1 protein (l, white bars) or on control plants (grey bars, see Material and Methods), show that AGL24 is able to bind to regulatory region IV (k, fragments 4 and 5) and SOC1 to regulatory region V (l, fragment 6). (m,n) Transient assays showing transactivation of the LUC reporter driven by region IV (activation by 35Sp:AGL24) and region V (activation by 35Sp:SOC1). 35Sp:NGA3 (NGA3 is an unrelated TF*(18)*) was used as an additional negative control.

**Fig. 3:**
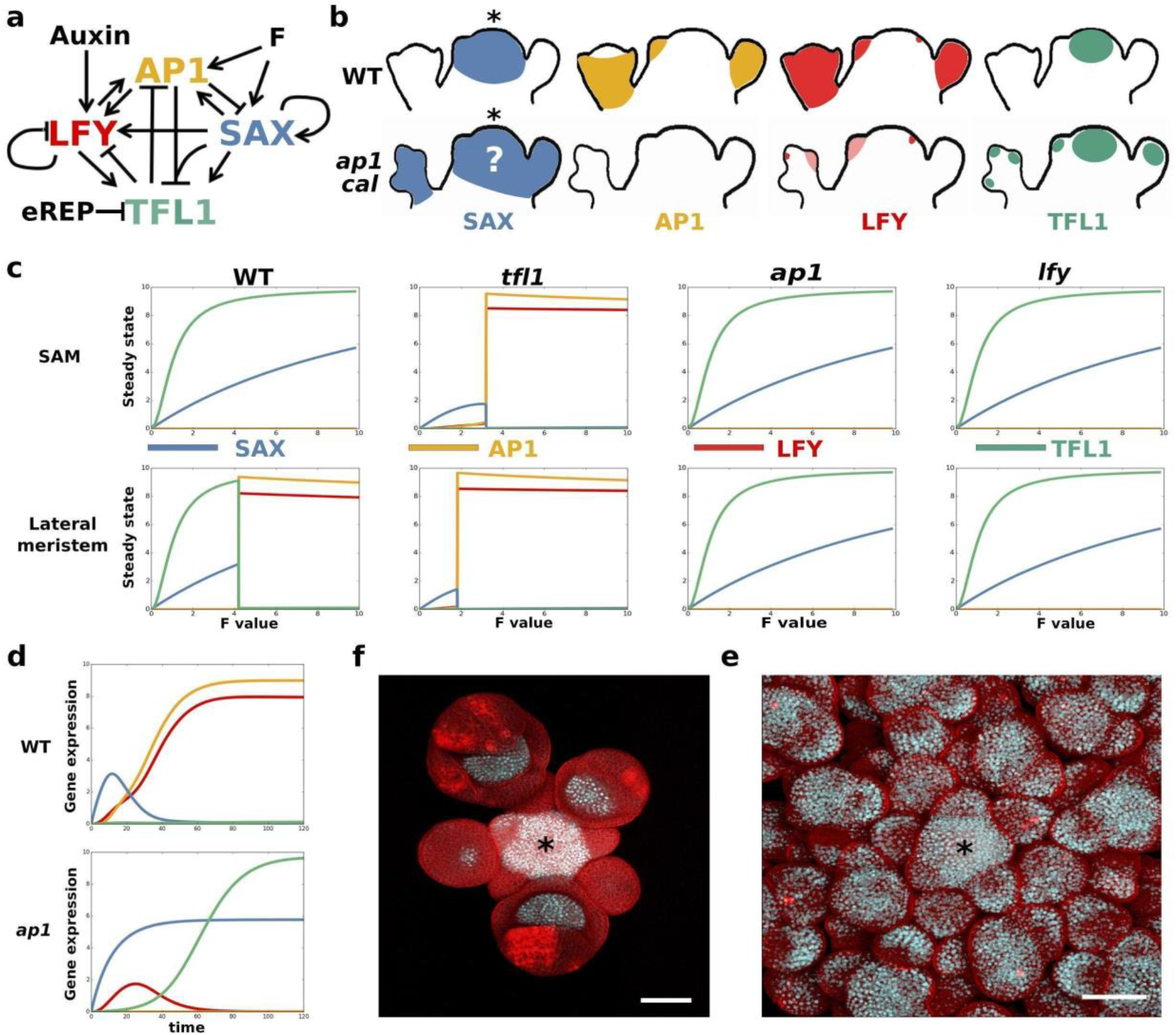
SALT GRN model and experimental validation. (a) SALT GRN network structure. Arrows correspond to positive regulations, and barred lines correspond to inhibitory interactions. (b) Known schematized expression patterns of ‘SAX’, ‘AP1’, LFY, and TFL1 in the SAM and lateral primordia of WT and *ap1 cal* mutant. ‘AP1’ and ‘SAX’ represent the approximated expression pattern of AP1 and CAL, and SOC1, AGL24 and XAL2, respectively. The question mark indicates a predicted expression pattern. (c) WT, *tfl1*, *ap1 cal* and *lfy* steady states of the model at different F values in the SAM (i.e., low auxin), and in lateral meristems (i.e., high auxin). The genetic identity predicted for WT and all mutant meristems correspond to the experimentally observed phenotypes. (d) Temporal simulation of gene expression in lateral primordia with high F value. In WT, a transient expression of SAX gives way to LFY and ‘AP1’ accumulation. In the *ap1 cal* mutant, the absence of ‘AP1’ activity prevents LFY stabilization and ‘SAX’ and TFL1 repression. (e, f) Expression of the SOC1:GFP (white/light blue signal) reporter construct in wild type background (e) and the *ap1-7 cal-1* mutant (f) flowering shoots. Asterisks mark the SAM. Scale bar = 50μm.

We thus created the SALT network (for SAX, ‘AP1’ for AP1/CAL, LFY and TFL1; Fig. 3a) made of these 4 regulators and their relationships, the auxin phytohormone *(20)*, and F, a flower inducing signal (a proxy for the FT florigen) that increases when the plant ages or is exposed to environmental conditions that favour flowering *(21,22)*. We also added a short-lived transient signal denoted eREP, a proxy for the *TFL1* early repression in the young flower bud performed by the SOC1/AGL24/SHORT VEGETATIVE PHASE/SEPALLATA4 redundant activity *(23)*.

The SALT network robustly generates the gene expressions observed in wild-type vegetative (low SALT values), inflorescence (high TFL1/SAX, low ‘AP1’/LFY) and flower (low TFL1/SAX, high ‘AP1’/LFY) meristems, dependent on F and auxin input values (Fig. 3b,c, Fig. S2). Above an F threshold value, the network is bistable generating either a flower or an inflorescence state. Simulations of *tfl1*, *lfy*, *ap1 cal* mutants produce expected outputs consistent with experimentally reported gene expressions *(13,24–26)*(Fig. 3b, c). The simulated *sax* mutant did not reach a floral state, consistent with the late flowering behaviour of the *soc1 agl24* double mutant *(27)*.

The modelled gene expression dynamics (Fig. 3d) illuminate the fundamental differences between the wild-type and the cauliflower meristems: in a wild-type flower primordium, F induces *SAX*. SAX and auxin induce *LFY*, that, together with F, induce ‘*AP1*’. ‘AP1’ positively feeds back on *LFY* and represses *SAX* (Fig. 3a). *TFL1* expression, that could potentially be induced by the SAX in early floral stages, is constantly repressed, first by eREP and later by SAX+‘AP1’. The high ‘AP1’ and LFY expressions together with low TFL1 and SAX levels stabilizes the floral fate. In contrast, in the *ap1 cal* flower primordia, the absence of ‘AP1’ activity has two consequences: i) *LFY* expression is upregulated only transiently since ‘AP1’ positive feedback is missing (Fig. 3d) and ii) *SAX* genes are not repressed by AP1 and thus induce *TFL1* in nascent flower meristems. TFL1 represses *LFY* even further and the meristem returns to a shoot meristem state (Fig. 3d). An obvious output of this model, not yet experimentally validated, is that *SAX* genes expression should extend over the entire cauliflower structure. To test this, we analysed a SOC1-GFP reporter line and indeed observed a striking expansion of its expression in *ap1 cal* as compared to the wild type (Fig. 3e, f).

The SALT network is thus able to recapitulate realistic gene expression in various mutant backgrounds. However, the global architecture of a plant does not solely depend on the genetic regulation of the different meristem fates: the *lfy* and *ap1/cal* backgrounds have the same steady state predictions (Fig. 3c) but very different plant architectures *(28)*. To solve this apparent discrepancy and try to understand the conspicuous cauliflower shape emergence, we integrated the SALT GRN in a realistic 3D model of plant development so that the plant architecture emerges as a property of a multitude of local GRN-based decision rules taken at the level of each meristem and of coordinated organ growth dynamics (Supplementary information).

## A multi-scale model generates Arabidopsis cauliflower structures

The 3D model is made of the 4 types of organs that shape plant architecture development: meristems, internodes, leaves and flowers (Fig. 4a). Each meristem has a molecular state computed at each instant by the GRN as a function of the meristem’s previous state and external factors, here auxin and F, themselves provided by the plant growth model. Meristem fates are determined by these states: low SAX, ‘AP1’, LFY and TFL1 values correspond to a vegetative identity that therefore produces a compressed stem (non-elongated internodes) making rosette leaves; high SAX and TFL1 coupled to low LFY and ‘AP1’ values confer a flowering shoot (inflorescence) fate where the meristem produces an elongating internode stem, a reduced cauline leaf and a new shoot meristem in the leaf axil; low SAX and TFL1 with high LFY and ‘AP1’ trigger the production of an internode stem terminating with a flower meristem. Such flower meristems are devoid of bracts (leaf-like organs) as their development is known to be repressed by LFY in Arabidopsis *(29)*. Each newly generated axillary meristem runs the GRN in its own state. Those meristems start with auxin level initially set to the maximum since they emerge in response to a local auxin accumulation *(20)*, and with SAX/LFY/‘AP1’ values inherited from the parent meristem, together with a fraction of the parent F and TFL1 value as these proteins diffuse from their synthesis regions (Supplementary information)*(21,30)*. To match the wild-type plant architecture, indeterminate meristems at orders >2 (see Fig. 4a for how meristem orders are defined) were kept quiescent, a likely effect of apical dominance (the inhibition of lateral branches growth by the primary SAM) (Fig. S3a). The model starts with a single apical vegetative meristem with low SAX, ‘AP1’, LFY, TFL1, F and auxin values. As the plant ages, the F increases, leading to newly produced lateral meristems with different identities. The model also contains laws describing the growth dynamics of the different organs (internode and leaves elongation, flowers growth, organ production rate, growth initiation delay (See supplementary materials).

**Fig. 4:**
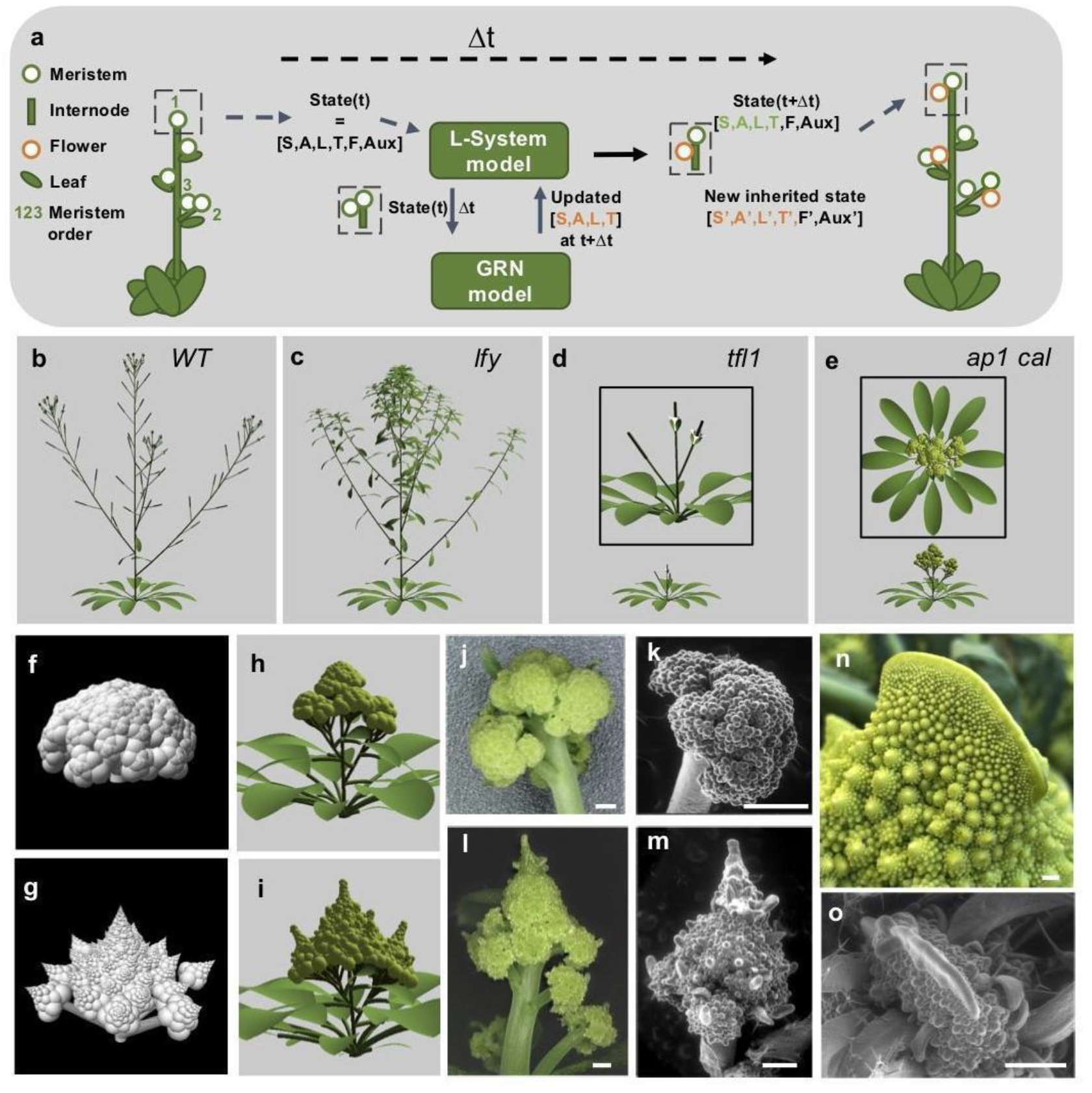
3D phenotypes with a GRN-based plant development model. (a) Schematic representation of the multi-scale model of Arabidopsis development. The model comprises four main organs: leaves, internodes, flowers (orange circles) and meristems (green circles). Each meristem has a proper state composed of signal levels (auxin, F) and GRN steady state. At any time *t*, the plant is made of a collection of organs (left). To compute the plant at time *t*+Δ*t* (right) the model updates the signal levels and GRN state in each meristem. The steady state defines the identity of the meristems (vegetative, inflorescence or flower) that are used to compute meristem lateral productions. Green numbers indicate meristem order (b-e) Plant morphologies obtained in the WT (b), *lfy* (c), *tfl1* (d) and *ap1 cal* (e) simulations. Simulated morphologies with constant (f,h) or increased meristem size (g,i) in a simplified (f,g) and the Arabidopsis model (h,i). Light micrographs (j, l,n) and s.e.m (k,m,o) of cauliflower structures in Arabidopsis *ap1 cal* (j, k), Arabidopsis *ap1 cal clv3* (l, m, o) and Romanesco (n). Uninduced *AP1:GR* transgene is present in plants j-m. Scale bar = 500 μm.

By adjusting the GRN and growth dynamics parameters within a range of realistic values (see Supplementary information), we calibrated the model dynamics to produce realistic architecture development for wild-type plants (Supplementary Movie 1), as well as for the *lfy* (Supplementary Movie 2) and *tfl1* mutants (Fig 4b-d) and a non-flowering phenotype for the *sax* mutant (not shown). However, in the simulation of *ap1 cal* mutants, the increased meristem reiterations observed in the real *ap1 cal* cauliflower morphology could not be obtained. Indeed the model does not include any gene function that instructs non-floral meristems of order > 2 to escape from arrest, as observed in real *ap1 cal* mutants (Fig. S3a-b), showing that the cauliflower phenotype involves additional regulations. To address this issue, we reasoned that laterally produced inflorescence meristems from *ap1 cal* are different from those produced in other genotypes as, according to our new GRN, they have been transiently exposed to LFY expression (Fig. 3d). We wondered whether this transient LFY expression could contribute to high order meristem release in addition to the bract suppression it already performs. Indeed, we have shown in a previous experiment that ectopic expression of LFY (or of a LFY allele) causes accelerated development of otherwise inhibited flower or shoot meristems on the main stem in the rosette *(31)* (Fig. S3i-k). Moreover, the *lfy ap1 cal* triple mutant does not form cauliflowers and we found that it has reduced high-order meristem development as compared to *ap1 cal* (Fig. S3d-h) supporting the hypothesis that *LFY* transient expression releases meristem growth on high-order axes.

We thus added to the model the condition that meristem arrest is released if the meristem is exposed early to a minimal threshold of LFY. Simulations showed that this is sufficient to unlock the recursive growth of lateral meristems and to generate the curd structure in the *ap1 cal* mutant, without any alteration of wild type growth dynamics (Fig. 4e, h, Supplementary Movie 3). The model thus shows that compact curds can emerge as a transient structure due to the massive and recursive production of meristems at all orders before internodes start to elongate. The cauliflower and *lfy* inflorescences are different despite having meristems in the same states (Fig. 3.c) because of the molecular histories that lead to these states are different. This phenomenon, called hysteresis, here derives from the specific bi-stable nature of the GRN controlling WT inflorescence development. In its altered *ap1/cal* version, GRN’s bistability is lost. However, what remains is a natural tendency to transiently maintain flower inducing cues such as LFY due a partial and irreversible incursion in the GRN flower program. Whether or not lateral meristems have been transiently exposed to LFY expression before is critical. No exposition will lead to the presence of bracts and prevent higher-order meristem development while a transient exposition induces bract suppression (leaving a single meristematic structure) and high-order lock release. Combined with the natural self-replication of the meristem *ad infinitum*, this produces the cauliflower phenotype without any other major modification of the wild-type growth dynamics, neither the meristem production rate nor the internode elongation speed.

## *Brassica oleracea* cauliflower and Romanesco curds structures can be simply explained by altered growth dynamics

Because *Arabidopsis thaliana* and *Brassica oleracea (Bo)* both belong to the Brassicaceae and their curds follow similar ontogenesis, we speculated that similar genetic processes could underlie both morphologies. Analysing RNA-seq data of *Bo* cauliflower curds, we confirmed the presence of a mutation in the *BoCAL* gene (Fig. S4a) and established that the two *AP1* paralogs (*BoAP1a* and *BoAP1c)* do not contain mutations in their coding sequence but are expressed at undetectable or extremely low levels in the curd (Fig. S4b). The *B. oleracea* cauliflower phenotype is thus likely due to the combination of mutations in the *BoCAL* coding sequence and mutations in *cis* or *trans* reducing *BoAP1* genes expression. At the morphological level, the overall structure of the cauliflower plant appears as a single massive curd explained by a number of meristem 56) much higher than in *B. oleracea* from which it originates (*Bo* var. *acephala*, maximal order is usually 2-3). The conical Romanesco curd shape (Fig. 1f), however, remains unexplained. Comparing the development of different types of *Brassica oleracea* curds, Kieffer and colleagues *(32)* identified several morphological parameters that remain constant during cauliflower development but vary in Romanesco. They are i) the plastochron, the time between two successive meristem productions, ii) the number of visual parastichies originating from a given meristem, iii) the number of plastochrons needed before a lateral primordium starts producing its own primordia (or lateral production onset delay), and iv) the size of the meristems. Whether some of these parameters are causal to the Romanesco phenotype has remained elusive.

Based on results from phyllotaxis studies *(1,33,34)*, it is established that the first three parameters are linked to the meristem size: an augmentation of the size of the meristem central zone should decrease the plastochron, which in turn increases the number of parastichies, and the lateral production onset delay. We thus hypothesized that passing from a constant to a decreasing plastochron in all plant meristems could change cauliflower into Romanesco morphologies. We first tested this *in silico* using a simplified, purely geometric model of curd growth, independent from the Arabidopsis genetic and morphological constraints (see Supplementary information). Strikingly, this single change was sufficient to produce conspicuous Romanesco shapes (Fig. 4g) whereas constant values of this parameter produces cauliflower morphologies (Fig. 4f).

We then introduced the same change in the more complex GRN-based, Arabidopsis cauliflower architectural model, while keeping its organ growth dynamics previously calibrated on the WT. Although not as perfectly as in the purely geometric model, the curd changed towards a “Romanesco-like” morphology with typical conical curd shapes (Fig. 4h, i). We then wished to test this hypothesis experimentally in the genetically amenable Arabidopsis plant by affecting the size of the meristem central zone directly. We achieved this by introducing a mutation in the *CLAVATA3* (*CLV3*) gene known to control meristem homeostasis and induce an increase of the meristem central zone during growth *(35–37)*. As predicted by our analysis the introduction of a *clv3* mutation in *ap1 cal* Arabidopsis mutant notably modified the curd shape that lost its round morphology for a more conical one, with similar structures at different scales, two features recognized as a hallmark of Romanesco curds *(38)* (Fig. 4l-m). Interestingly, the *ap1 cal clv3* triple mutant also occasionally developed extreme fasciation caused by dramatic meristem enlargement, a feature also commonly observed in Romanesco curds (Fig. 4n, o). Altogether, these observations establish that meristem size is a fundamental regulator of the final curd morphology, notably through the control of plastochron value.

Our results provide evidence that the self-similar fractal nature of cauliflower curd spiral patterns is primarily due to specific perturbations of a floral GRN that prevent meristems to completely switch on the flower development program. However, these floral incursions are not totally reversible and leave a trace in the corresponding meristems that partially acquire flower attributes due to their early exposure to LFY (leaf repression and insensitivity to apical dominance). Combined with the impossibility to repress the inflorescence identity genes (SAX and TFL1) and to release higher-order meristem growth, this confers to those meristems the ability to iterate again and again attempts of making flowers everywhere and up to high orders, leading to an explosion of meristems activity. Internode elongation being less rapid than organ initiation, this process progressively accumulates a compact, and extremely self-similar organ, in the form of a fractal sphere-like curd, before it unfolds in the adult plant when meristem activity markedly decreases. If this initiation process accelerates during organogenesis (due to meristem size drift), the curd produces conical structures instead of spherical ones, that visually reveal, through the myriad of self-nested spirals, the striking fractal nature of the curd by repeatedly deploying conical structures in 3-dimensions, such as in Romanesco curds.

## Acknowledgments

We thank Anne-Marie Chèvre, Teva Vernoux, Chloe Zubieta and Hicham Chahtane for advice, Dominique Tardy, Eric Giraud, Renaud Dumas and Vincent Martin (OBS, France) for providing cauliflower samples, Richard Immink (Wageningen, Netherlands), C. Ferrándiz (IBMCP; Spain), George Coupland (MPIPZ, Germany), Miguel Ángel Blázquez (IBMCP, Spain), Richard Amasino (UWM, USA) and the European Arabidopsis Stock Centre for providing seeds, Vincent Berger (CEA/DRF) for the Keyence microscope, Christine Lancelon-Pin (Plateau de microscopie électronique - ICMG. CERMAV-CNRS) for SEM experiments.

## Author contributions

ChG and FP conceived the study

ChG, EA, EF performed the modelling

ASM, CaG, DB, FM, FP, GT, MK, MLM, VG designed and performed the plant experiments

NP performed the confocal imaging experiment

JL analysed the RNA-seq data

ChG, FP and EA wrote the paper with the help of all authors

## Supplementary Materials

Materials and Methods

Figures S1-S6

Tables S1-S3

Movies S1-S3

Supplementary References

