## Supplemental_materials for "The making of cauliflowers: the story of unsuccessful flowers"

### The fractal making of cauliflowers: the story of an unsuccessful flower

**This PDF file includes:**

Materials and Methods
Supplementary Text
Figs. S1 to S6
Tables S1 to S3

**Other Supplementary Materials for this manuscript include the following:**

Movies S1 to S3

**MATERIALS AND METHODS**

### Plant materials

The following *A. thaliana* lines in the Columbia-0 background were used: *lfy-12 (1)*; *ap1-7 (2), cal-1 (3), ap1-7 cal-1* double mutants (gift from J. Goodrich, Institute of Molecular Plant Science, University of Edinburgh, Edinburgh, United Kingdom), SOC1p-GFP *(4)* (provided by D. Posé), *35Sp:FT (5)*, *35Sp:SOC1 (6)*, *soc1-2 (6)*, *AGL24p:AGL24-RFP (7)* and *agl24-2 (8)*. *35Sp:CO (9)*, *co-3 (10)*, *ft-3 (5)*, *35S:AP1-GR ap1-1 cal-1 (11)* and *TFL1p:GUS* (this work) *are A. thaliana* lines in the Landsberg *erecta* background, as well as the *35S:AP1-GR ap1-1 cal-1 (11)* used for the mutation of *CLV3* using the CRISPR-Cas9 strategy.
For crosses between accessions, it was previously verified that the GUS reporter activity was not affected in wild-type Col x L*er* hybrids.
*Brassica oleracea* var. botrytis and Romanesco cauliflowers were obtained from Observatoire Breton des Semences (OBS) or bought at the Grenoble local market.

### Plant growth conditions

Seeds were sown on soil or surface-sterilized and grown in Petri dishes on Murashige and Skoog basal salt mixture medium (Sigma-Aldrich, www.sigmaaldrich.com). Seeds were stratified 1-3 days at 4°C, transferred mixture of phagnum:perlite:vermiculite (2:1:1) or on soil:vermiculite (5:1) and grown at 21-22°C under long-day (16 h) or short-day conditions (8 h). *ap1 cal* plants were grown at 18°C for 16 h light and 16°C for 8 h dark, under 65% humidity. *Nicotiana benthamiana* plants for transient assays were grown on a mixture of sphagnum:vermiculite (1:1) under long-day photoperiod at 24°C.

### Plasmids construction

For the reporter analysis of *TFL1* expression, we produced a *TFL1p:GUS* construct that contained the *β-glucuronidase* (*GUS*) gene flanked by 2177 bp of the *TFL1* promoter 5' region and 4605 bp of the *TFL1* promoter 3' region *(12)*. The 5' region was amplified from a L*er* genomic clone with primers 5'-TGA<u>GTCGAC</u>GCTAGGAGACTTCGTTGATC-3' (*Sal*I site is underlined) and 5'-CTGCA<u>GGATCC</u>TTTTCTTTTGTTAACTTAGAGG-3' (including the ATG of the *TFL1* gene, *BamH*I site is underlined). This fragment was cloned as *Sal*I-*BamH*I into the pBI101 vector *(13)*, upstream the *GUS* gene. The 3' region was amplified with primers 5'-GTAGAGCTCTAGATTCATGATTGTCATAAACTGC-3' (including the stop codon of *TFL1*, *Sac*I and *Xba*I sites are underlined) and 5'-TAGAATTCGGTACCAAGTTGAAGTCTCTCATTGACGAAC-3' (*Eco*RI and *Kpn*I sites are underlined). This fragment was cloned as *Sac*I-*Eco*RI into the pBI101 vector, thus replacing the nopaline synthase terminator downstream the GUS gene. The resulting *TFL1p:GUS* cassette was cloned as *Sal*I-*Kpn*I into the pBIN19 binary vector. This construct was introduced into wild-type Arabidopsis plants of the L*er* ecotype by vacuum infiltration *(14,15)*. An homozygous line carrying the transgene in a single locus was selected for further analysis.

For transient expression assays, the cDNAs of *SOC1* and *AGL24* were cloned into the pMDC32 vector by Gateway LR recombination to create *35Sp:SOC1* and *35Sp:AGL24* plasmids respectively. The *35Sp:NGA3* plasmid was already available *(16)*. The *TFL1* promoter region IV (fragment between +2823 and + 3230 bp downstream the *TFL1* stop codon) was amplified with forward primer 5'-CTCGAGGACTCTCGAGGACAAACCAAC-3' (*Xho*I site is underlined) and reverse primer 5'-CTTATATAGAGGAAGGGTCTTGATTATGGGTTAGCTATAAAGATGG-3' (overlapping sequence with a 35SminΩ promoter is underlined). The *TFL1* promoter region V (fragment between +3420 and +3752 bp downstream the *TFL1* stop codon) was amplified with forward primer 5´-CTCGAGCGGATTGGTCCAGTTAGAAC-3' (*Xho*I site is underlined) and reverse primer 5'-CTTATATAGAGGAAGGGTCTTGAAGAAGCTCCTACCACTTGAAG-3' (overlapping sequence with a 35Smin *(17)* promoter is underlined). A -90 bp CaMV 35S promoter that includes the omega translational enhancer (35S )*(17)* was amplified with forward primer 5'-CAAGACCCTTCCTCTATATAAG-3' and reverse primer 5'-CCATGGTGTAATTGTAAATAGTAATTGTAATGTTG-3' (*Nco*I site is underlined). The 35SminΩ sequence was fused by PCR downstream the *TFL1* promoter regions with the described region-specific forward primers and the 35SminΩ reverse primer. The resulting fragments were cloned as *Xho*I/*Nco*I into the pGreenII 0800-LUC vector *(18)*.

For the CRISPR Cas9 mediated mutation of *CLV3*, three gRNA spacers specific of the *CLV3* gene were designed using CHOPCHOP https://chopchop.rc.fas.harvard.edu/ and synthetized by eurofins. Guide sequences are the following:

- Guide1: 5'-GATTGGAGACCAGAAGCATCATGA-3'

- Guide2: 5'-GATTGGTTTCTTGGCTGTCTTGGT-3'
- Guide3: 5'-GATTGGTGAATGGGTTGGAGCAAA-3'

These guides were used to build one vector with two gRNA spacers (guides 1-2) and one with the three gRNA (guides 1-2-3)

After annealing oligonucleotide guides, each of the 3 double-stranded DNAs were ligated in pBSK-AtU6-26:guide vector *(19)*, modified to allow sequential ligation of several guides *(20)*. Resulting plasmids were verified by sequencing. *Kpn*I/*Sbf*I restriction fragments containing the assembled guides and the UBQ10:CoCas9:tUBQ10 cassette of the pBSK:CoCas9 digested by *Sbf*I and *Eco*RI were simultaneously ligated in the pCAMBIA plant transformation vector containing the At2S3:eGFP as selection marker. This plasmid was obtained by cloning the At2S3:eGFP:t35S reporter construct from pFP100 *(21)* into *Bst*X1/*Psp*X1 digested pCAMBIA1300 backbone. The pBSK:CoCas9 contains a codon-optimized version optimized for expression in Arabidopsis of the human Cas9 under the control of the UBQ10 promoter *(20)*. Final plasmids (pMLM17 with 2 guides and pMLM18 with 3 guides) were verified by sequencing.

### *Arabidopsis thaliana* transformation

*Agrobacterium tumefaciens*, strain C58C1pMP90, carrying pMLM17 or 18 were grown overnight at 28 °C in Luria Broth medium supplemented with rifampicin 50 µg/mL, gentamycin 50 µg/mL and kanamycin 50 µg/mL and used for the Arabidopsis floral dip method *(22)* using Silwet L-77 at 0.01 %. To improve transformation efficiency of the *35S:AP1-GR ap1-1 cal-1* plants, a spray with 10 mM Dexamethasone supplemented with Silwet L-77 0.01 % was performed 10 days before the agrobacterium floral dip to trigger AP1-GR nuclear translocation and induce flower development and seed recovery.

### Molecular characterization of *CLV3* CRISPR-Cas9 lines

The effect of *clv3* mutation in *ap1 cal* were studied by transforming MLM17 and MLM18 constructs in *35S:AP1-GR ap1-1 cal-1*. T1 primary transformed seeds were collected based on their seed fluorescence under a SZX12 Olympus dissecting microscope.

We obtained 14 T1 plants with pMLM17 vector (5 out of 14 showing pyramidal curds) and 16 with the pMLM18 (7 out of 16 showing pyramidal curds). PCR amplification of the *CLV3* genomic sequence around the sgRNA target sites was used to detect Cas9/sgRNA induced mutations and deletions. gDNA was extracted from leaves using Edwards extraction buffer (200 mM Tris-HCl pH7.5 ; 250 mM NaCl, 25 mM EDTA, 0.5 % SDS 20 %). For the PCR amplification, two couples of primers were used oMLM1134/oMLM1131 (166 bp) and oMLM1132/oMLM1133 (268 bp)

oMLM1131: 5'-CATGAGCTTGAGTGAGATCTGG-3'
oMLM1132: 5'-AATGTTGTTCAATTGGCAGATG-3'
oMLM1133: 5'-CCGAAATGGTAAAACCGATAAA-3'
oMLM1134: 5'-GCTACTACTACTACTCTTCTGC-3'

In the T2 generation, non-fluorescent seeds were sown to select against the transgene. PCR product obtained with oMLM133/1134 primers were sequenced to check mutations at the *CLV3* locus. Sequencing revealed several types of mutations in the *CLV3* gene. Plant MLM18-25-4 has a T nucleotide insertion at position 57 creating a frame-shift with a stop codon in the 1$^{st}$ exon ((First exon sequence with added t and bold stop codon: ATGGATTCGAAGAGTTTTCTGCTACTACTACTACTCTTCTGCTTCTTGTTCCTTCAt**TGA**) amino acid sequence: MDSKSFLLLLLLFCFLFL**H**), as well as a deletion of 100 bp between guides 3 and 2. The mutant MLM17-3 shows the same insertion of a T nucleotide as the MLM18-25-4 mutant. Plants shown in figure 4 are MLM18-25-4 T3 (Fig. 4j), MLM17-3 T1 (Fig. 4k) and MLM18-4-28 T3 (Fig. 4m).

To verify that the pyramidal phenotype observed in *ap1 cal* 35S:AP1-GR is due to a single mutation, one T2 plant (MLM18-2-19 in *35S:AP1-GR ap1-1 cal-1*) was crossed to *35S:AP1-GR ap1-1 cal-1* and F2 progenies from two individual F1 plants were analyzed. The pyramidal phenotype was observed in 6 out of 35 (17%) and 8 out of 40 (20%) F2 plants consistent with a 25% / 75% segregation with a single mutation at the *CLV3* locus (Chi-square test of independence p=0.205 > 0.5).

As a control, pMLM18 construct was transformed into the Col-0 and L*er* wild type backgrounds. 27 and 30 T1 plants were obtained in Col-0 and L*er* backgrounds respectively, all showing typical *clv* phenotype including increased petal number, increased pistil size due to increased carpel number and inflorescence meristem size.

### Seed and plant imaging

For selection of MLM17 and MLM18 transgenics based on At2S3: GFP fluorescence in seeds, an Olympus SZX12 dissecting microscope with fluorescent illumination was used. Light micrographs were taken with the Keyence digital microscope (VHX-5000) using a Z20 X50 lens.

Confocal imaging was performed as described *(23)*. Environmental scanning electron microscopy experiments were performed at the Electron Microscopy facility of the ICMG Nanobio-Chemistry Platform (Grenoble, France). Untreated flowers were directly placed in the microscope chamber. Secondary electron images were recorded with a Quanta FEG 250 (LV) microscope while maintaining the tissue at 2°C, under a pressure of 120 Pa and a 25 % relative humidity. The accelerating voltage was 7 kV and the image magnification ranged from 100 to 800X.

**GUS assay and histological procedures**

Arabidopsis shoot apices were stained for GUS as described *(12)* with minor modifications. Staining buffer contained 10 mM of potassium ferri- and ferrocyanide in order to minimize leakage of the X-Gluc reaction product *(24)*. Photographs of whole-mount samples were taken with a Color View 12 digital camera connected to a Nikon SMZ800 binocular stereoscopic microscope. After GUS staining, some samples were cleared with cloral hydrate solution for 2-3 days at 4°C *(25)*. Images of cleared apices were taken with a Nikon DS-Fi1 digital camera connected to a Nikon Eclipse 600 light microscope under Nomarski interference optics. For tissue sectioning, samples were stained in 0.2% (w/v) eosin and embedded in paraffin as described *(12)*. 12 μm sections were obtained with a Leica RM-2025 microtome and images were taken under bright-field microscopy.

**Luciferase transient assays**

4-week-old Nicotiana plants (with 5-6 leaves) were used for the analysis. Agrobacterium cultures (C58pMP90 strain) were incubated for 24 hours at 28°C (to stationary phase, $OD_{600}$ 1-2). Cells were collected and resuspended in infiltration buffer (10 mM $MgCl_2$, 10 mM MES and 0.2 mM acetosyringone) to an $OD_{600}$ of 0.4. Agrobacterium mixtures with a 1:5 reporter:effector ratio were prepared. These were incubated for 3 hours at room temperature and darkness with gentle shaking (50 rpm) before infiltration of young and fully expanded Nicotiana leaves. Three days later, firefly luciferase (LUC) and renilla (REN) activities were assayed from 0.5 cm leaf discs (approximately 20 mg) using the Dual-Luciferase® Reporter Assay System kit according to the manufacturer's instructions (Promega). Absolute relative luminiscence units were measured with a

GloMax™ 96 Microplate Luminometer (Promega). LUC/REN ratios were averaged from at least three biological replicates (different plants) each one with three technical replicates (different leaves from the same plant).

**Chromatin immunoprecipitation assay**

The commercial antibody Living Colors_DsRed polyclonal antibody (Clontech) and an antibody against a synthetic peptide (SVKCIRARKTQVFK) were used for AGL24 and SOC1 ChIP experiments respectively. Chromatin was prepared from inflorescences and at least three independent experiments were done. Wild-type (Col-0) plants were used for chromatin extraction for the SOC1 ChIP experiment, and *soc1-2* plants served as a negative control. Homozygous *AGL24p:AGL24-RFP agl24-2* plants were used for the AGL24 ChIP experiment and wild-type plants served as a negative control. The ChIP assays were performed as described previously *(26)*. Six primer sets (fragments 1-6) were designed in the 3' *TFL1* promoter region: the forward primer 5'-AAACGTGGAGATACGGAAAAGG-3' and the reverse primer 5'-ACCAGCCGTGAAAATAGATATG-3' for fragment 1, the forward primer 5'-GCATTCTACATTGATTCAGTG-3' and the reverse primer 5'-TGAATTAATGACACGTGACC-3' for fragment 2, the forward primer 5'-GTTTTAGGGTTTCAGTAACAC-3' and the reverse primer 5'-ATGGAATGGAACAGAGCACG-3' for fragment 3, the forward primer 5'-GGTCCAAGGGTTAGTATGTTTC-3' and the reverse primer 5'-GCCGCAAACCTGGTGATTAACC-3' for fragment 4, the forward primer 5'-GAAACATACTAACCCTTGGACC-3' and the reverse primer 5'-GTAAATGTACCTCCTCGTCAC-3' for fragment 5, and the forward primer 5'-TCAATTTCGGATTGGTCCAG-3' and the reverse primer 5'-CTTAGTTGTAACTGATGAACG-3' for fragment 6. Enrichment folds were detected by qPCR using a SYBR Green assay (Bio-Rad). The assay was performed in triplicate using a Bio-Rad C1000 Thermal Cycler optical system and relative enrichment was calculated as previously described *(26)*.

**Cauliflower curd sequencing**

Curds from developing cauliflowers grown in the field (planted July 10th 2017, harvested Oct 18-27 2017) were provided by O.B.S (Breton Seed Observatory): OBS_5045_1 (named OBS-Caul-1 on figure S5), OBS_800_1 (OBS-Caul-2), OBS-0795_2 (OBS-Caul-3), OBS_800_2

(OBS-Caul-4), OBS_0795_1 (OBS-Caul-5), OBS_0819_1 (OBS-Caul-5). We also used two cauliflowers (Samples CAUL_1, CAUL_2a, CAUL_2b) and two Romanesco (Samples ROM_1, ROM_2a, ROM_2b) bought on the Grenoble producer market in 2017. 2a and 2b correspond to extractions made on different parts of the same curd. Total RNAs were extracted using the RNeasy Plant Kit protocol (QIAGEN) followed by DNAse treatment and sent to Genewiz (UK) for quality control, Poly-A selection, library cloning and sequencing (Illumina HiSeq4000, 2x150bp configuration, single index).

The obtained reads were aligned with the HDEM Reference Genome *(27)* with STAR (outFilterMultimapNmax 1; outFilterMismatchNmax 6; alignIntronMax 30000; other parameters set by default). FeatureCount (default settings) was used to detect reads mapping to *Brassica oleracea* genes. EdgeR (default settings) was then used to identify the genes differentially expressed between the individuals analyzed. The orthologs of *AP1* (At1g69120), *CAL* (At1g26310) and *FUL (*At5g60910) of cabbage are identified by BLAST protein against the *Arabidopsis thaliana* proteome (TAIR10). The Heatmap of the expression of these genes is generated by a python ad hoc script.

The following secure token has been created to allow review of record GSE150627 while it remains in private status: khkjgckmdtkhpgb.

### MODELING METHODS

#### Regulatory network construction and simulation

#### ALT network

The SALT model included the genes SAX genes (i.e., *SUPPRESSOR OF OVEREXPRESSION OF CONSTANS 1* (*SOC1*), *AGAMOUS-LIKE 24* (*AGL24*) and *XANTAL2* (*XAL2*)), *APETALA1* (*AP1*) and *CAULIFLOWER* (*CAL*) (hereinafter *AP1*), *LEAFY* (*LFY*), and *TERMINAL FLOWER1* (*TFL1*). These genes are the key transcription factors (TF) that have been experimentally associated with the flowering transition and the development of cauliflower structures in *A. thaliana*. We also included two inputs, auxin, a phytohormone that promotes the initiation of new primordia, and F that represents different types of molecules inducing the flowering phase. Because we did not include the auxin signaling pathway and F represents multiple flowering inducing pathways, the activity of both auxin and F is only phenomenological in our model. Auxin concentration is maximal in newly formed primordia, while its activity is basal in the shoot apical meristem. For this reason, shoot apical meristems and newly formed primordia states are different in their auxin levels. As the value of F increases, we expect the shoot apical meristem to transit to an inflorescence state, and to a flower state for lateral primordia. Based on the experimental information listed in Sup Table 1, we build the GRN described by the next equations:

$$\frac{dS}{dt}=\left(\frac{k_{A,S}^{n}}{k_{A,S}^{n}+A^{n}}\right)\left(\frac{F}{k_{F,S}+F}+\frac{S^{n}}{k_{S,S}^{n}+S^{n}}\right)-\left(k_{a_{S+A}}SA-k_{d_D}D\right)-\delta S$$

$$\frac{dA}{dt}=\left(\frac{k_{T,A}^{n}}{k_{T,A}^{n}+T^{n}}\right)\left(\frac{L^{n}}{k_{L,A}^{n}+L^{n}}+\frac{S^{n}}{k_{S,A}^{n}+S^{n}}\frac{F}{k_{F,A}+F}\right)-\left(k_{a_{S+A}}SA-k_{d_D}D\right)-\delta A$$

$$\frac{dL}{dt}=\left(\frac{k_{T,L}^{n}}{k_{T,L}^{n}+T^{n}}\right)\left(\frac{aux+aux_b}{k_{aux,L}+aux+aux_b}\frac{k_{L,L}^{n}}{k_{L,L}^{n}+L^{n}}\frac{S^{n}}{k_{S,L}^{n}+S^{n}}+\frac{A^{n}}{k_{A,L}^{n}+A^{n}}\right)-\delta L$$

$$\frac{dT}{dt}=\left(\frac{S^{n}}{k_{S,T}^{n}+S^{n}}\frac{k_{R,T}^{n}}{k_{R,T}^{n}+R^{n}}\frac{k_{D,T}^{n}}{k_{D,T}^{n}+D^{n}}\right)+\left(\frac{L^{n}}{k_{L,T}^{n}+L^{n}}\right)-\delta T$$

$$\frac{dD}{dt}=k_{a_{S+A}}SA-k_{d_D}D-\delta D$$

$$\frac{daux}{dt}=-\delta aux$$

$$\frac{dR}{dt}=-\delta R$$

*S*, *A*, *L*, *T* and *aux* represent *SAX, AP1*, *LFY*, *TFL1*, and auxin respectively. *D* and *R* represent the dimer *SAX-AP1*, and the eREP repressor of *TFL1*. $aux_b$ represent a basal level of auxin activity. $k_{X,Y}$represents the binding affinity of variable $X, X=\{S,A,L,T,D,R,F,aux\}$, to the promoter region of gene $Y, Y=\{S,A,L,T\}$. $k_{a_{S+A}}SA$ and $k_{d_D}$ are the association rate of AP1 with SAX and the dissociation rate of the SAX-AP1 complex. We assumed that the degradation $\delta$ of all variables were equal to *0.1* minutes, which is a realistic value given that the half-life of proteins is usually within the hours range *(28)*. Finally, the level of cooperativity $n$ of variable $X$ for inducing gene $Y$ for all X and Y genes were equal to 2, representing the formation of protein dimers for gene regulations, which is common for gene regulation *(29)*.

We tried to optimize the SALT model randomly choosing different sets of values of $k_{X,Y}$ within sensitive biological values (see Parameters below).

**Parameters**

Data for the exact value of the parameters included in the network are not available in the literature. However, we used general knowledge about the different processes to constrain the parameter space. In particular we considered that the concentration of TFs in cells is within the nanoMolar (nM) range *(30)*. The mean life time of proteins, including TFs is within the hours range *(28)*. The affinity of transcription factor to the DNA is usually lower than $10^{-8}$M (i.e., affinity≤μM)*(31)*. Finally, TFs, including TFL1, AP1, and LFY bind to hundreds of target genes, and all TFs can randomly bind to the DNA *(32)*. Thus, the effective concentration of TFs is lower than the real concentration. Assuming that the effective concentrations of the TFs is 2 orders of magnitude smaller than the real concentration, we constrain $k_{X,Y}$ values within the range [$10^{-10}$M, $10^{-6}$M].

The SALT model was able to produce the expected behavior, and the kinetic was optimized by hand (see Table S2). Then, we analyzed the effect of modifying the value of the $k_{X,Y}$s. To do this, we fixed the value of all $k_{X,Y}$ to its optimal value, except one, that we varied within the interval [$10^{-10}$M, $10^{-7}$M]. As observed in Fig. S2. The overall behavior of the network was maintained for most $k_{X,Y}$ values.

**Structure of the model**

Further the previous section, the model structure (as reported in Fig. S6, with feedback loops and feedforward loops (FFL) listed in Table S3), allows for some analysis which is independent of specific parameter values. The "mutual repression" positive feedback listed in Table S3 is solely mediated by AP1. The other positive 2-loop, between AP1 and LFY, is visible but cannot affect the competition with TFL1: it plays a role of stabilisation, or self-maintenance, of the floral identity. One consequence of this loop is that, at steady state, LFY and AP1 are always expected to be either both high or low. It is apparent, without having to write any detailed model of the dynamics, that the removal of AP1 above deprives the network from its ability to overcome TFL1's expression; even if external signals transiently increase LFY's expression, without AP1's contribution this cannot be maintained permanently and the system must reverse to a ``non-flower'' identity (high TFL1). The structure of the positive three loop entails that any steady state must not only have AP1 and LFY at similar levels, but that this should be opposite to TFL1 level, again confirming the antagonism floral/non-floral identities.

Negative feedback loops are typically expected to confer homeostasis, and potentially lead to oscillatory behaviour if they comprise at least three elements. Since there is no evidence of oscillations of this system, the negative 3-loop mentioned above is likely to not have a functional role, but appear as a byproduct of the overall structure of the network. The LFY-TFL1 negative loop has the potential to drive both variables to intermediary values, effectively maintaining the system at the interface between longer periods of time than would occur with positive feedback loops only.

In broad terms, *(71)* characterizes coherent FFLs to induce a response delay (to “off steps” for type 2 and “on steps” for type 4, given the AND nature of these FFLs), which can have the function of a “persistence detector” (i.e. their last term responds to pulses of the first term only if it persists over time), while incoherent FFLs have the potential to speed up response times (the “on step” for type 1 and “off step” for type 3, with also a potential to induce pulses of TFL1 if LFY is present).

Combining the feedforward and feedback loops leads to the following interpretation: the GRN comprises a core “ALT” network, which has an intrinsic ability to present clearly distinct floral (high LFY and AP1, low TFL1) and non-floral attractors (high TFL1, low AP1 and LFY), while also presenting homeostatic properties able to transiently maintain the network in intermediary states (all three core variables at intermediate levels). On top of this core network, a layer of feedforward loops is able to speed up the triggering of the core ALT network’s internal dynamics, while protecting the flower inducing genes from transient removal of the florigen signal. Without the AP1 node, the remaining LEAFY-TFL1 “core” network is irremediably deprived of its bistability, and only high TFL1/low LFY can be achieved as a permanent state. However, two remaining FFLs (6th and 11th in Table S3) have the potential to make the increase (in response to SAX, i.e. FT after a delay) of LFY faster than that of TFL1 and make the removal of TFL1 (upon SAX removal) delayed compared to that of LFY. Both are consistent with the occurrence of a transient pulse of LFY upon FT induction via SAX, during which TFL1 remains low. This analysis indicates that the model robustness to parameter changes is an intrinsic consequence of its structure and would persist beyond the parameter exploration mentioned above.

**Coupling of plant architecture model and GRN**

To model the interaction between the GRN studied in this work and the development of plant architecture, we developed a model for organ and branching system development.

We coupled the developmental model with the GRN model presented in the previous section.

Our developmental model consists of a set of developmental rules expressing the growth of the plant apices and the elongation of internodes. These rules are encoded using the L-system formalism (33) in the L-Py programming language *(34)*.

At any moment, the plant is represented as a set of components organized in a bracketed string and called the L-string. Components within a pair of matching brackets correspond to the successive components of a given axis. The position of an opening bracket in the L-string defines the position of the axis in the tree branching system. Components themselves represent the various plant components (meristems, internode, leaves, etc.) that can be found in a plant branching system. Each component can bear attributes describing its state (e.g. diameter, length, genetic state, etc.). Altogether, the L-string represents the current state of the plant branching system. This string is made of three main types of components denoted $I$ (internodes), $A$ (apices), $L$ (leaves).

For example, the L-string:

L = $I(5{,}21)I(5{,}19)[L(10{,}40)][I(3{,}10)A(s_1)]I(4{,}12)[A(s_2)]A(S_0)$

represents a plant made of 3 axes:

1. $I(5{,}21)I(5{,}19)[L(10{,}40)]I(4{,}12)A(s_0)$
2. $I(3{,}10)A(s_1)$
3. $A(s_2)$

In this example, internodes $I$ have two attributes, namely $d$ and $l$, corresponding respectively to their average diameter and length. Likewise, leaves $L$ have two attributes corresponding to their maximum width and length. Apices $A$ have a single vector attribute aggregating several variables defining the state of an apex. $A$ typical such state contains variables corresponding to plastochron information, gene expression and signal levels.

Let's assume for instance that the state of each apex is governed by a GRN composed of 3 genes, $A, B, C$. Then the state of every apex will contain a value corresponding to the level of expression of this gene in the apex state. In our example, each state $s_i$ of apex $i$ will contain the following variables: $s_i.A$, $s_i.B$, $s_i.C$, corresponding to the expression levels of genes in state $s_i$, and a plastochron count down value $s_i.PCD$, counting the time remaining before initiating a new lateral organ.

Based on these definitions, assuming a constant increment of time $\Delta t$ between the simulation steps, the rules of development can be written as rewriting rules of the form:

$I(d,l)$:

compute $dl$ as a function of the increment of time $\Delta t$

**produce** $I(d, l + dl)$

$L(\omega,l)$:

compute $dl$ as a function of the increment of time $\Delta t$

compute $d\omega$ as a function of the increment of time $\Delta t$

**produce** $L(\omega + d\omega, l + dl)$:

$A(s)$:

compute the new state $ns$ as a function of the old state $s$ and $\Delta t$

**if** $s.PCD == 0$:

compute an initial state $is$ for the new lateral apex as a function of the old state $s$

**produce** $[A(is)]A(ns)$

**if** $s.PCD == 1$:

**produce** $A(ns)$

The first two rules define how each internode and leaf $L$ are growing with time. Increment of the geometric variables (length, width and diameter) characterizing these components are computed over a time lapse $\Delta t$, and are used to update the values of internode and leaf geometric variables. The time evolution of an apex is slightly more complex. First the new state $ns$ is computed based on the apex previous state $s$ and on $\Delta t$. This updates the value of the gene expression levels contained in the apex current state (detailed below). Then, if the plastochron countdown has reached 0 ($s.PCD == 0$:), it triggers the production of a new lateral apex with an initial state partially inherited from the main apex's state (see below), and an apical state with updated state. Otherwise, one just replaces the current apex with an updated one.

Finally, a set of interpretation rules make it possible to map at each time point the current L-string to a detailed geometric interpretation extended from Mundermann et al. (35).

These rules are essentially cosmetic and their detailed description can be found in the available code.

### Integration of the GRN model

At time step $t$, each apex $i$ is in a state $s_i$ stored as an attribute of the corresponding component $A(s_i)$ in the L-string. We assume that this state is characterized by some attractor of the SALT GRN described in section Regulatory network construction and simulation. During development, this state can change if one of its entries, aux or F, is forced to a new value by the architectural processes implemented in the L-systems rules and external to the GRN *per se*.

Let us denote $g_1, \dots, g_k$ the set of GRN variables and call $g_1, \dots, g_{k_0}$ subset of the entries that might be overwritten by external processes during the plant growth (variables that can be forced from the outside in the GRN). At the end of the $n^{th}$ time step, the state of an apex $i$ is stable, defined by $s_i$, and the $k^{th}$ gene having the value $s_i.g_k$. During the next step of duration $\Delta t$, apex processes or aging may change the value of one or several of the input gene variables $g_1, \dots, g_{k_0}$. In this case, the apex GRN state from the previous state may no longer be valid, i.e. compatible with the GRN model. Therefore the GRN transition function must be called with the new GRN input gene values to compute the attractor corresponding to this new situation.

The previous rule related to apex growth must thus be modified as follows:

$A(s)$:

define an empty structure $ns$ for recording the new state

evaluate the input variables $g_1, \dots, g_{k_0}$ as a function of the old state $s$ and of $\Delta t$

compute the new GRN attractor given the new values of $g_1, \dots, g_{k_0}$

update the new state $ns$ with the new computed GRN attractor

finalize the computation of the new state $ns$ as a function of the old state $s$ and $\Delta t$

**if** $s.PCD == 0$:

compute an initial state $is$ for the new lateral apex as a function of the old state $s$

**produce** $[A(is)]A(ns)$

**if** $s.PCD == 1$:

**produce** $A(ns)$

Between two simulated steps, the state of every apices is thus evaluated using this rule. The state of newly initiated lateral meristems is derived from the apical meristem state, using a set of putative inheritance rules. In particular, a new lateral apex inherits a fraction of TFL1 and SAX level from the apical meristem, while LFY and AP1 expressions are set equal to zero. Thanks to this tight coupling between plant architecture model and GRN, the architecture of the plant can be seen as an emerging property of the two-way interaction between GRN and growth. It made it possible to reproduce all the mutant phenotypes related to genes involved in the cauliflower GRN.

**Geometric model of Romanesco**

To study the different cauliflower morphologies, we used the L-system model, following the rules described in the previous section. Then, we defined a constant and a variable plastochron countdown. For the constant countdown:

$$s.PSD(t+\Delta t) = s.PSD(t+\Delta t) - cte$$

while for the variable countdown:

$$s.PSD(t+\Delta t) = s.PSD(t+\Delta t) - (cte)(s.AGE)$$

where $s.AGE$ is the age of the meristem since it was created.

**Supplementary figures**

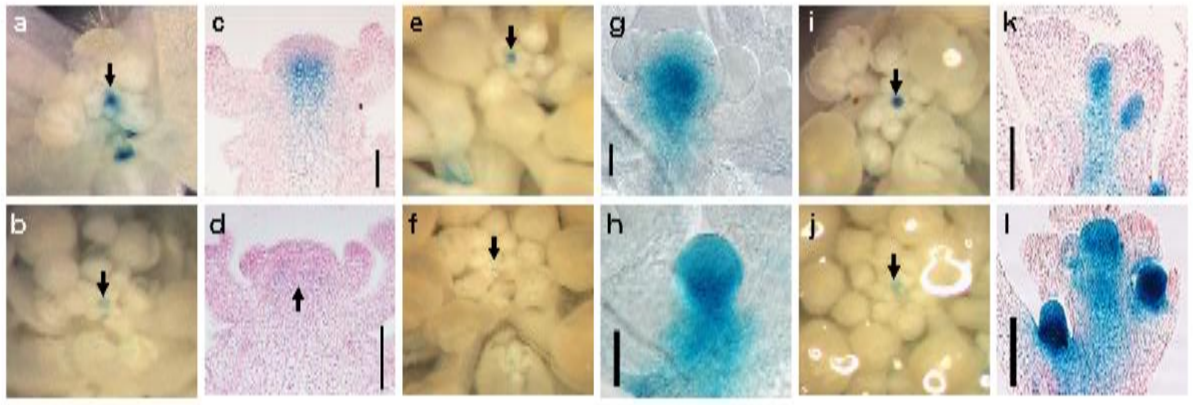


**Figure S1. The photoperiod flowering pathway upregulates TFL1p:GUS activity in the shoot apex. (a-b)**, TFL1p:GUS activity in representative shoot apices grown under long-day (a) or short-day (b) photoperiod. **(c-d)**, longitudinal sections through the apices shown on the left. **(e-f)**, TFL1p:GUS activity in representative WT (e) and *co-3* (f) shoot apices at a similar developmental stage. **(g-h)**, TFL1p:GUS activity in cleared WT (g) and *35Sp:CO* (h) shoot apices. **(i-j)**, TFL1p:GUS activity in representative WT (i) and *ft-3* (j) shoot apices at a similar developmental stage. **(k-l)**, TFL1p:GUS activity in longitudinal sections through representative WT (k) and *35Sp:FT* (l) shoot apices at a similar developmental stage. Arrows mark the SAM region. Scale bars in (c), (d), (k) and (l), 100 µm. Scale bars in (g) and (h), 40 µm.

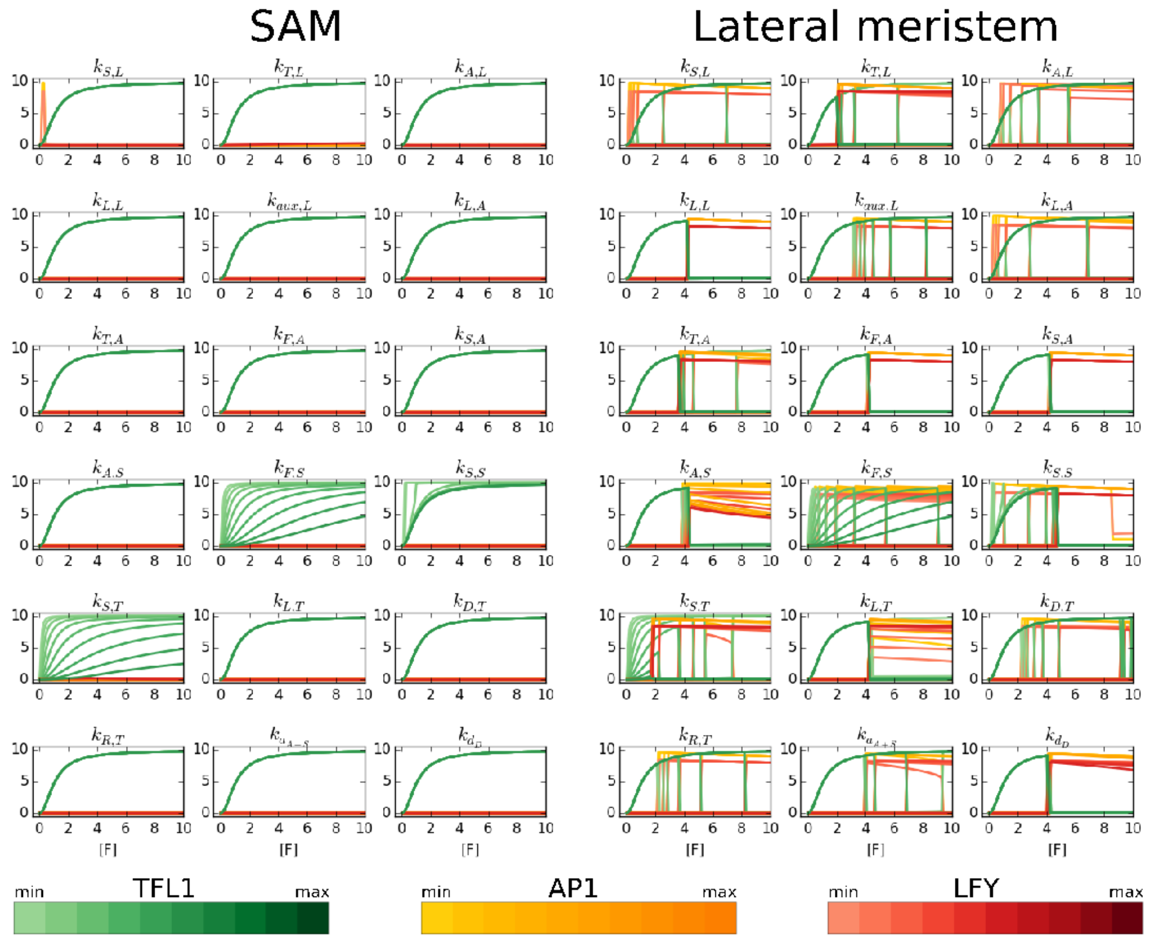


**Figure S2: Robustness of the SALT model**

Steady state of the SALT GRN (y axis) at different F values (x axis) using 10 different values of the parameter indicated above each graph (see Modeling Methods section), while the other parameters remain fixed. The first three columns of graphs show the analysis in SAM conditions (i.e., without auxin), and the next three columns show the analysis performed in lateral meristem conditions (i.e., with auxin). The value of the parameters changes in regular intervals from 0.001 (light green, yellow and red colored curves) to double its optimized value (dark green, yellow and red colored curves). Green, yellow and red curves correspond to TFL1, AP1, and LFY value, respectively. The same qualitative behavior is observed independently of changes in the value of the parameters.

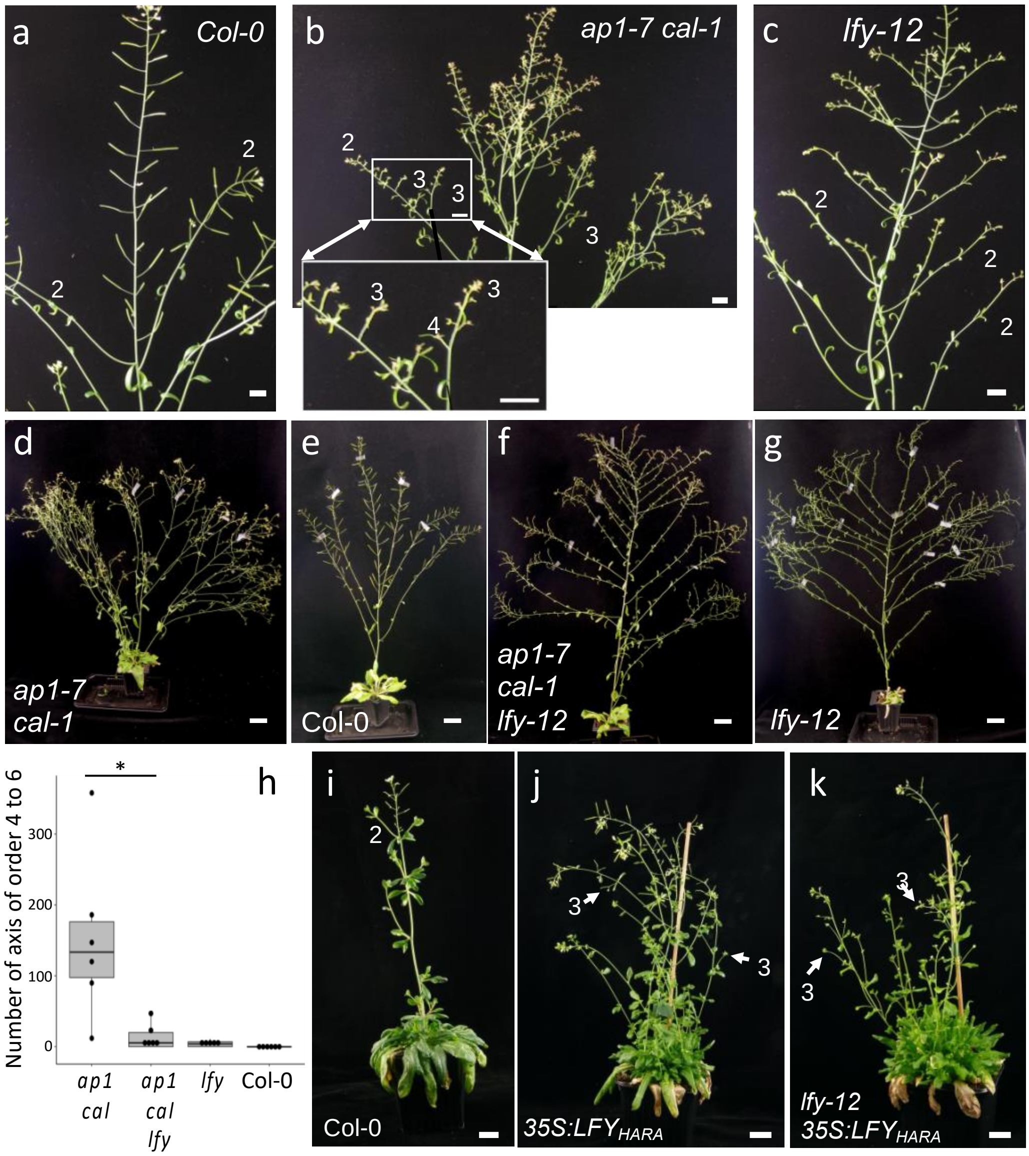


**Figure S3: Genetic control of lateral axis development:** Lateral axes of higher order develop in *ap1 cal* , in a *LFY* dependent manner, compared to wild type or *lfy* plants.

1ary inflorescences are order 1 inflorescences, 2ary inflorescences (or coflorescences) developing on 1ary inflorescences are order 2 inflorescences, and so on... Numbers on the pictures below indicate the order of the closest inflorescence.

(a-c) Inflorescences of six-week-old plants from grown in long day. Whereas in this growth conditions, inflorescences of orders >2 are very rarely observed in Col-0 (a) or *lfy* mutant (c), several of them are visible in *ap1 cal* mutant (b). Scale bar = 1 cm. (d-f) The highly branched structure of *ap1 cal (d)* compared to wild type Col-0 (e) is lost in *ap1 cal lfy* triple mutant (f), that have an architecture very close to the *lfy* single mutant (g). Plants were grown for 2 months in long days conditions. Scale bar = 5 cm. Scoring axis of order

4 and higher per plant (h) confirms that the *ap1 cal lfy* plants behaves differently from *ap1 cal* plants (Wilcoxon test, * p-value 0.0099) (i-k) Two-month-old short day grown plants ectopically expressing of a modified version of LFY (35S:$LFY_{HARA}$) in Col-0 (j) or in *lfy-12* (k) backgrounds[1] show order 3 axes that are not observed in wild type plants (i), suggesting that the exposure to $LFY_{HARA}$ triggers their development, beside triggering rosette leaves meristem development as demonstrated in Chahtane et al.[1]. Scale bar = 2 cm

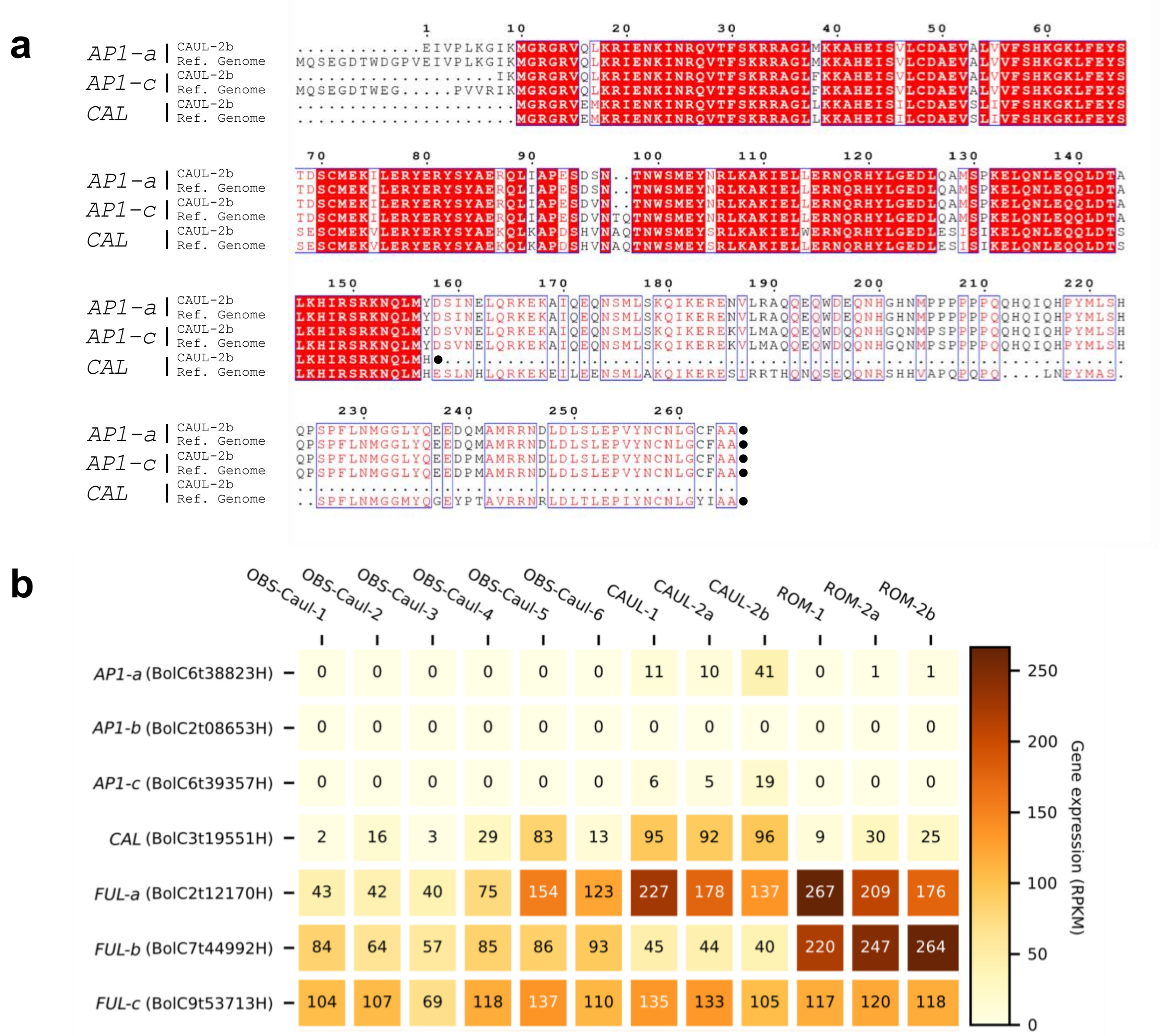


**Figure S4: Sequence and expression of *AP1* and *CAL* genes in different cauliflower curds.** (a) Sequence alignment of the AP1-a, AP1-c and CAL proteins from *Brassica oleracea*. Sequences of the proteins were translated from the *Brassica oleracea* reference genome *(27)* or from the gene sequence as assembled from reads obtained from RNA-seq of cauliflower curds (sample CAUL-2b). (b) Expression level of *AP1*, *CAL* and their close *FRUITFUL* (*FUL)* homologs based on RNA-seq. For each gene, the normalized number of reads (RPKM) is shown as a value and color coded according to the color scale. OBS are cauliflowers harvested in field of the Brittany Seed Observatory, CAUL and ROM samples are issued from two mature curds bought on the market (2a and 2b are different samples from curd 2). *AP1* genes expression is undetectable in growing cauliflowers (OBS samples) and weak in mature cauliflowers as compared to *FUL* genes, the closest *AP1* homologs.

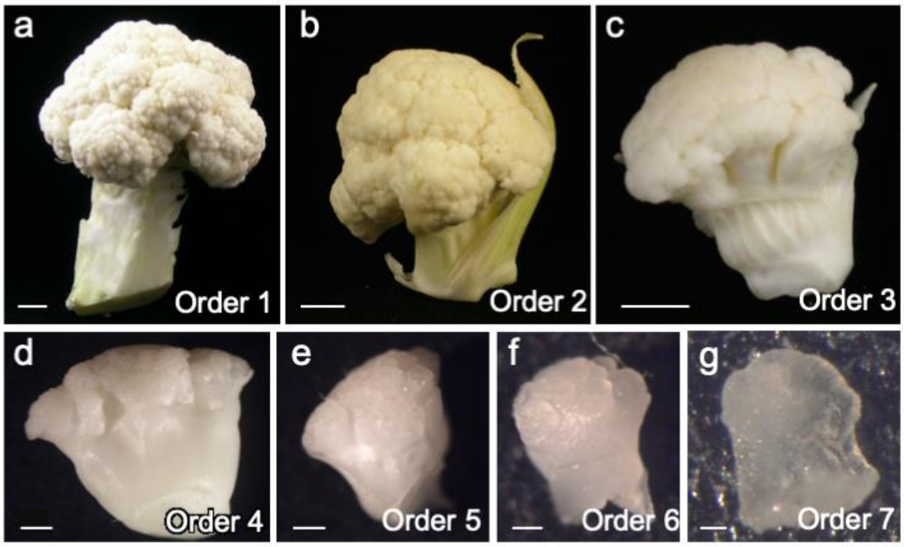


**Figure S5**: **Dissection of a young cauliflower curd** (a) to reveal axes of successive orders from order 2 to 7 (b-g). Scale bars: 2 cm (a), 0.5 cm (b, c), 1 mm (d), 500 µm (e), 200 µm (f), 50 µm (g)

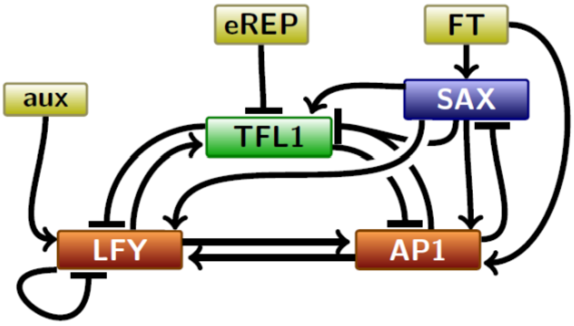


**Figure S6**: **Structure of the GRN model.** The nodes represent the main variables included in the model. Arrows represent their regulations (→ for activation and ⊣ for repression).

**Table S1. List of interactions used in the GRN.**

| Regulator | Type of regulation | References |
|---|---|---|
| ***AP1/CAL* regulations** | | |
| **LFY** | **Positive direct** | (36–40) |
| **TFL1** | **Negative. Probably indirect** | (41–44) |
| **SAX** | **Positive direct** | *(3,45)* |
| **F** | **Direct in complex with FD** | (46–50) |
| ***LFY* regulations** | | |
| **AP1/CAL** | **Positive direct** | *(2,51,52)* |
| **TFL1** | **Negative direct** | (42–44) |
| **SAX** | **Positive direct** | (3,53–56) |
| **Auxin** | **Positive via MP/ARF5** | *(57)* |
| **LFY** | **Negative direct** | (36–38,58,59) |
| ***TFL1* regulations** | | |
| **AP1/CAL** | **Negative direct** | *(2,51,52,60)* |
| **SAX** | **Positive direct** | This study, *(45)* |
| **SAX** | **Negative direct** | *(45,61)* |
| **LFY** | **Positive direct** | (36–38,62,63) |
| **CAL** | **Negative** | *(43)* |
| ***SAX* regulations** | | |
| **AP1** | **Negative direct** | *(45,52,64,65)* |
| **SAX** | **Positive direct** | *(4,8,45,53,56,66)* |
| **F** | **Positive direct** | (54,55,67–70) |

**Table S2. Parameter values used in the SALT GRN.**
All $k_{X,Y}$ are in nM t[-1] units. Time (t) is in arbitrary units.

| | |
|---|---|
| $k_{L,A}$ | 1 |
| $k_{T,A}$ | 1 |
| $k_{F,A}$ | 50 |
| $k_{S,A}$ | 15 |
| $k_{A,S}$ | 4 |
| $k_{F,S}$ | 10 |
| $k_{S,S}$ | 20 |
| $k_{S,T}$ | 1 |
| $k_{L,T}$ | 100 |
| $k_{D,T}$ | 1 |
| $k_{R,T}$ | 0.1 |
| $k_{S,L}$ | 6 |
| $k_{A,L}$ | 4 |
| $k_{L,L}$ | 100 |
| $k_{T,L}$ | 5 |
| $k_{aux,L}$ | 1 |
| $k_{a_{S+A}}$ | 0.1 |
| $k_{d_D}$ | 0.01 |
| $\delta$ | 0.1 |
| $n$ | 2 |

**Table S3. Feedback and feedforward loops in the SALT GRN.** Based on Figure S6.

| Feedback loops and their sign | Coherent and Incoherent FFLs (ignoring SAX) |
|---|---|
| ▪ Mutual activation AP1←→LFY (>0)<br>▪ Mutual repression AP1⊢⊣TFL1 (>0)<br>▪ Positive 3-loop LFY→AP1⊣TFL1⊣LFY (>0)<br>▪ Self repression LFY ⊢⊣LFY (<0)<br>▪ Negative 2-loop LFY⊢→TFL1 (<0)<br>▪ Negative 3-loop LFY→TFL1⊣AP1→LFY (<0) | ▪ CFFL, type 2: TFL1⊣AP1→LFY / TFL1⊣LFY<br>▪ CFFL, type 2: TFL1⊣LFY→AP1/ TFL1⊣AP1<br>▪ CFFL, type 4: AP1⊣TFL1⊣LFY/ AP1→LFY<br>▪ IFFL, type 3: AP1→LFY→TFL1/ AP1⊣TFL1<br>▪ IFFL, type 1: LFY→TFL1⊣AP1/ LFY→AP1<br>▪ IFFL, type 1: LFY→AP1⊣TFL1/ LFY→TFL1<br>▪ SAX->LFY->AP1 / SAX->AP1; CFFL type 1<br>▪ SAX->LFY->TLF1 / SAX->TFL1; CFFL type 1<br>▪ SAX->AP1->LFY/ SAX->LFY; CFFL type 1<br>▪ SAX->AP1-\|TLF1/ SAX->TFL1; IFFL type 1<br>▪ SAX->TFL1-\|LFY / SAX->LFY; IFFL type 1<br>▪ SAX->TFL1-\|AP1 / SAX->AP1; IFFL type 1 |
